# CNS antigen-specific control of early B-lineage development in the meninges

**DOI:** 10.1101/2021.06.03.446985

**Authors:** Yan Wang, Dianyu Chen, Di Xu, Chao Huang, Danyang He, Heping Xu

## Abstract

The random V(D)J recombination of immunoglobulins (Ig) loci often creates autoreactive B cell progenitors expressing self-recognized B-cell receptors (BCRs)^1^, which are eliminated or inactivated through an autoantigen-dependent central tolerance checkpoint to prevent autoimmune reactions^2,3^, a process thought to be restricted to the bone marrow (BM) in the adult mammals^4^. Here we report that early developing B cells are also present in the meninges of mice at all ages. Single cell RNA-sequencing (scRNA-seq) analysis revealed a consecutive trajectory of meningeal developing B cells in mice and non-human primates (NHPs). Parabiosis together with lineage tracing of hematopoietic stem cells (HSCs) showed that meningeal developing B cells are continuously replenished from the HSC-derived progenitors via a circulation-independent route. Importantly, autoreactive immature B cells which recognize myelin oligodendrocyte glycoprotein (MOG)^5^, a central nervous system (CNS)-specific antigen, are eliminated from the meninges but not BM. Furthermore, genetic deletion of MOG restored the self-reactive B cells in the meninges. Thus, we propose that meninges function as a unique reservoir for B cell development, allowing *in situ* negative selection of CNS-antigen-autoreactive B cells to ensure a local non-self-reactive immune repertoire.

## Main

Brain is long believed to be an immune privileged organ with inadequacy to mount an efficient immune response due to lack of lymphatic drainage and unreachability of immune cells^6^. However, this concept has recently been revised by the (re)discovery of a meningeal lymphatic system, which allows immune cells to reach the CNS under homeostatic condition^7,8^. For instance, single-cell profiles have revealed that B cells are present in the meninges of mouse brain^9,10^. On the other side, it’s known that autoreactive developing B cells can escape the tolerance checkpoints and persist in the mature B cell repertoire, when local concentration of tissue-specific self-antigen is insufficient to initiate the negative selection in the BM and spleen^5,11,12^. In this context, self-reactive B cells that recognize highly CNS-specific antigens could also escape the negative selection and reach the meninges through the meningeal lymphatics. Therefore, whether and how autoreactive B cells respond to CNS-specific antigens in the meninges remains to be elucidated.

### B cell progenitors present in the meninges of mice

Through intravenous injection of APC-conjugated CD45 antibody to label and exclude circulating immune cells followed by intracardiac perfusion (**Fig.1a, Extended Data Fig. 1a**), we found that B cells (CD19^+^B220^+^) were present in the dura mater of adult mice at homeostasis (**Fig. 1b**). Whole-mount immunolabeling and imaging revealed that most of the B cells aligned linearly along Lyve1-expressing lymphatic vessels adjacent to the sinuses (**Fig. 1c, Extended Data Fig. 1b**). Unexpectedly, a large proportion of meningeal B cells lacked membrane expression of IgD and IgM (**Fig. 1d**), and expressed relatively low amount of CD45 and B220 (**Extended Data Fig. 1c**), mirroring those observed in the BM (**Fig. 1e, Extended Data Fig. 1d**). This is in striking contrast to other tissues including spleen, lymph node (LN), lung and small intestine (SI), in which essentially IgD^+^ cells dominated the B cell population. Notably, IgD^-^IgM^-^ B cell compartment was absent in the circulating B cell compartment that were labeled with intravenously injected CD45 antibody (**Extended Data Fig. 1a, 1e**), further highlighting the unique profile of meningeal B cells.

**Fig. 1:**
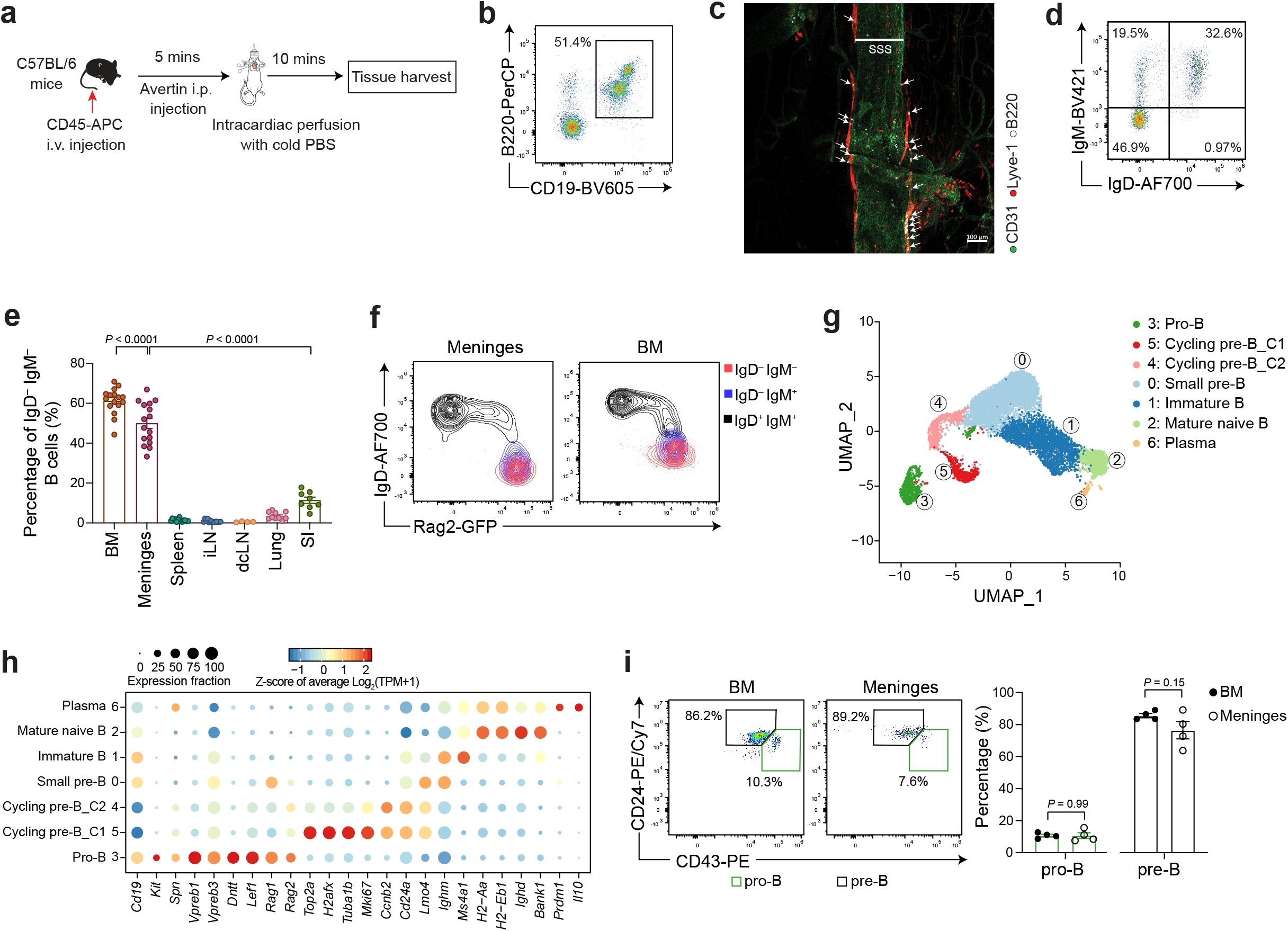
Identification of early B-lineage development in the meninges of mice. **a**, Experimental scheme. WT C57BL/6 mice were intravenously injected with CD45-APC antibody followed by anesthetization and intracardiac perfusion. Meninges, BM, spleen, LN, Lung and SI were harvested and subjected to tissue dissociation and immune cell isolation. **b**, A representative flow plot of B cells (CD19^+^B220^+^) in CD45^int/+^CD11b^-^ cell compartment (**Extended Data Fig. 1a**) in the dura mater of adult WT B6 mice. **c**, Representative image of CD31, Lyve-1 and B220 expressing cells around the superior sagittal sinus (SSS) region (as highlighted in **Extended Data Fig. 1b**). Arrows, B cells resided inside the lymphatics. **d**, A representative flow plot showing membrane IgD and IgM expression in meningeal B cells. **e**, Quantification of percentages of IgD^-^IgM^-^ cells in B cell (B220^+^CD19^+^) compartment of different tissues as indicated. **f**, Representative FACS plots showing Rag2-GFP expression in IgD^-^IgM^-^ (red), IgD^-^IgM^+^ (blue) and IgD^+^IgM^+^ (black) B cells isolated from the meninges and BM. **g**, Uniform Manifold Approximation and Projection (UMAP) learned two-dimensional (2D) representation of meningeal B cell profiles (dots), colored and numbered by cluster membership. Clusters are rank ordered by size. **h**, For representative differentially expressed genes (columns) across clusters (rows) in (g), shown is the fraction of cells in the cluster that express a gene (dot size) and the z-score of the mean expression of that gene in the cluster (color; z-score of average log2(TPM+1)). The inferred identities were labeled at the left. **i**, Proportions of pro-B (CD43^+^CD24^+^) and pre-B (CD43^-^CD24^high^) cells in IgD^-^ IgM^-^ B cell compartment in the BM and meninges. Data are representative of three biological replicates (b-d, f) or the pool from six biological replicates (g, h) or three independent experiments (e, i) shown as the mean ± s.e.m. Each symbol represents an individual mouse (e, i). Statistical significance was tested by one-way ANOVA followed by Tukey’s multiple comparisons test (e) or two-tailed t-test (i).

These observations promoted us to ask whether a proportion of meningeal B cells are at the early development stages, including progenitor (pro-) and precursor (pre-) B cells, both lacking membrane expression of IgM and IgD (IgD^-^IgM^-^), and immature B cells which express IgM but not IgD (IgM^+^IgD^-^). To determine the developmental stages of meningeal B cells, we employed an endogenous recombination-activating gene 2 (*Rag2*) reporter (RAG2-FP) mouse line^13^, in which only early lymphocyte progenitors are labeled with GFP. Indeed, similar to the expression dynamic in the BM, almost all IgD^-^ B cells in the meninges highly expressed Rag2-GFP (**Fig. 1f**), supporting an early developing stage. In contrast, no B cell expressing high amount of Rag2-GFP in other tissues was observed (**Extended Data Fig. 2a**). Notably, the transcripts of *Il7* and *Cxcl12*, which are essential niche genes supporting B cell progenitor development, were also readily detected in the meninges (**Extended Data Fig. 2b**). Three-dimensional imaging analysis of intact skull adhered with dura matter^14^ **(Extended Data Fig. 2c, supplementary movie 1**) revealed that Rag2-GFP-expressing B cells tended to align linearly within Lyve1-expressing lymphatic vessels in the meninges **(Extended Data Fig. 2d, supplementary movie 2**), while their counterpart in the skull mainly resided inside the bone marrow cavities at the interparietal area **(Extended Data Fig. 2e, supplementary movie 3**). Taken together, these results demonstrated that the meninges contain early developing B cells which were aligning closely to lymphatics.

To further characterize the transcriptional states of B cells in the meninges, we analyzed *Cd19*-expressing cells from the single cell RNA-seq dataset of CD45^+^CD11b^-^ cell population isolated from the meninges at neonatal (postnatal day 2 (P2)), postnatal (P14) and adult (P56) stages (**Online Methods**). We observed a consecutive development trajectory of B cell lineage composed by 7 different clusters (**Fig. 1g,h, Supplementary Table 1**), including pro-B (cluster 3, high expression of *Kit, Spn, Vpreb1, Dntt*, and *Rag1*), cycling pre-B (clusters 5 and 4, upregulated expression of *Mki67* and other mitotic genes but downregulated expression of *Rag1*) (**Extended Data Fig. 3a**), small pre-B (cluster 0, downregulated expression of mitotic genes^15^ but re-upregulated expression of *Rag1*), immature B (cluster 1, high expression of *Ighm* and *Ms4a1* but not *Ighd*), mature naïve B (cluster 2, high expression of both *Ighm* and *Ighd*) and plasma cells (cluster 6, high expression of *Prdm1* and *Il10*), arranged sequentially as B cell development occurs in the bone marrow^4^, and this B cell development trajectory is further recapitulated by the pseudotime analysis (**Extended Data Fig. 3b**). We validated the presence of pro-B (CD24^int^CD43^hi)^ and pre-B (CD24^hi^CD43^int^) cells in the meninges by flow cytometry, and found the frequencies of these cells in the meninges and BM were comparable (**Fig. 1i**). Whereas plasma cells were only present at adulthood when peripheral immune system have matured, the meningeal developing B cells were robustly detected from neonates to adulthood (**Extended Data Fig. 3c, d**), suggesting that the emergence of B cell progenitors in the meninges is not driven by the immaturity of peripheral immune system. In fact, the appearance of B cell progenitors (IgD^-^IgM^-^) persist in aged brain, albeit at a lower frequency (**Extended Data Fig. 3e, f**). This indicates the mechanism underlying the appearance of B cell progenitors in the meninges is distinct from those found in the gut laminar propria, which transiently appear at the age of weaning and decrease to undetectable levels by postnatal day 35^16^. Intriguingly, neither scRNA-seq (**Fig. 1g, h**) nor flow cytometry (**Extended Data Fig. 3g, h**) analyses detected any earlier progenitors, including pre-pro B cells and common lymphoid progenitors (CLPs), in the meninges, suggesting that meninges might function as a unique reservoir for B cell development but not early hematopoiesis.

### A consecutive trajectory of developing B cells in the meninges of NHPs

We next explored whether such early developing B cells also exist in the meninges of higher organisms, such as non-human primates (NHPs). To this end, we harvested dura mater and spleen of *M. mulatta* after intracardiac perfusion (**Fig. 2a**) for flow cytometry and scRNA-seq analysis (**Fig. 2b-d**). Notably, a group of cells with intermediate expression of CD45 (CD45^int^) was identified by flow cytometry in the dura mater, but not in the spleen or peripheral blood mononuclear cells (PBMCs) of NHPs (**Extended Data Fig. 4a**), mirroring what we observed in murine meninges that also contained CD11b^-^CD45^int^ fraction, a population enriching developing B cells (IgD^-^) (**Extended Data Fig. 1c, 4b**). As lacking a reliable antibody to label NHP B cells, we flow-sorted CD45^+^CD11b^-^CD3^-^ cells for scRNA-seq analysis to ensure sufficient representation of B cells in our dataset (**Fig. 2b**). Overall, nine clusters of B-lineage cells expressing distinct marker genes (**Supplementary Table 2**) were identified from the NHP meninges, exhibiting more heterogeneity than meningeal B cells of mice (**Fig. 1g**). For example, B cell clusters that expressed high amount of *Itgax* (cluster 1), *Cd1c* (cluster 3), *Il16* (cluster 5) and *Bcl6* (cluster 4) were not detected in the scRNA-seq dataset of mice meninges (**Fig. 1g, 2c**). Nevertheless, leveraging the expression dynamics of RAG, mitotic and other forementioned marker genes of early developing B cells, we were able to reliably annotate clusters of B cells at different developing stages, including pro-B (cluster 7), cycling pre-B (clusters 8), small pre-B (clusters 2) and immature B (clusters 0) (**Fig. 2d**). Importantly, the emergence of B cell progenitors is specific to the meninges, evident by the observation that none of splenic B cell clusters (**Extended Data Fig. 4c, Supplementary Table 3**) in NHPs manifested early developing identities, according to the expression of the same set of marker genes (**Extended Data Fig. 4d**). Taken together, these results demonstrated the presence of a consecutive trajectory of B cell development, starting from pro-B cell stage, in the meninges of both mice and NHPs under homeostatic condition.

**Fig. 2:**
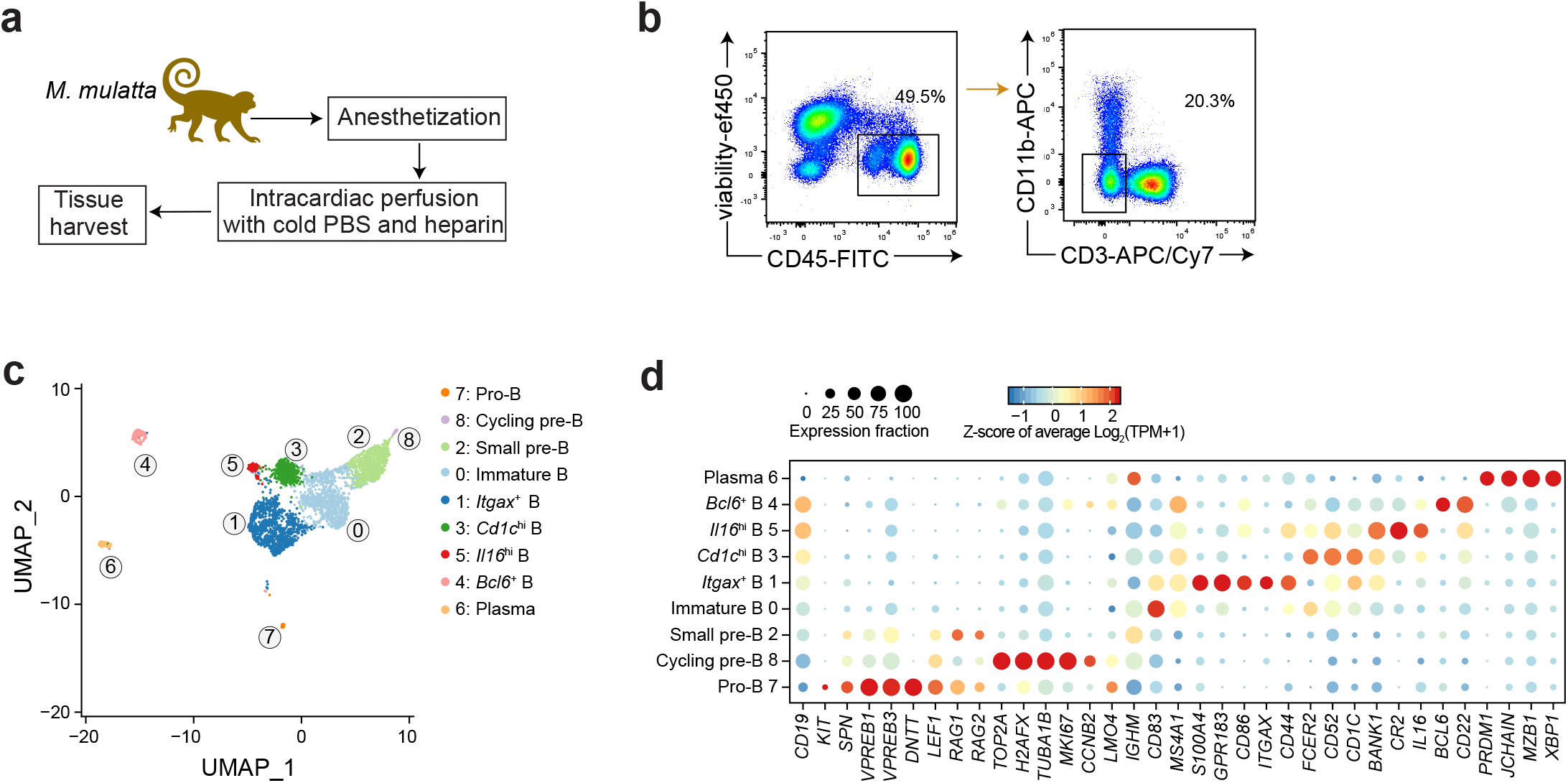
scRNA-seq analysis uncovers a consecutive trajectory of meningeal developing B cells in NHPs. **a**, Experimental scheme. After anesthetization and perfusion, the dura matter and spleen of *M. mulatta* were harvested and digested for immune cell isolation. **b**, Representative FACS plots for sorting CD45^int/+^CD11b^-^CD3^-^ cells. **c**, UMAP learned 2D representation of meningeal B cell profiles (dots), colored and numbered by cluster membership. Clusters are rank ordered by size. **d**, For representative differentially expressed genes (columns) across clusters (rows) in (c), shown is the fraction of cells in the cluster that express a gene (dot size) and the z-score of the mean expression of that gene in the cluster (color; z-score of average log2(TPM+1)). The inferred identities were labeled at the left. Data are representative (b) or the pool (c,d) of two replicates.

### HSC-derived progenitors continuously replenish meningeal developing B cells via a circulation-independent route

Given our findings of persistent existence of developing B cells but lack of CLPs and earlier progenitors in the meninges, we asked whether meningeal developing B cell population is constantly replenished by hematopoietic stem cells (HSCs)-derived progenitors from the BM. To test this, we traced the progeny cells differentiated from HSCs using a fate-mapping mouse model that expresses tamoxifen-inducible Cre-ERT recombinase under the control of the stem cell enhancer of the stem cell leukemia (Scl) locus (HSC-SCL-CreERT2)^17^ and a ROSA26-LoxP-STOP-LoxP-GFP allele^18^, in which HSCs and their progeny will be genetically labeled with GFP upon tamoxifen administration (**Fig. 3a**). We observed GFP-expressing cells among total CD19^+^ B cell population (~10%) in the meninges at two weeks post tamoxifen injection (**Fig. 3b**), a stage when the proportion of GFP^+^ B cells in the spleen, LN and lung remained low (<1%) (**Extended Data Fig. 5a**). These GFP^+^ CD19^+^ B cells in the meninges were mainly at the early developmental stages, evident by low expression of B220 and lack of IgD (data not shown), suggesting an emergence of B cell progenitors from HSCs in the meninges. Importantly, the frequencies of GFP-expressing B cells gradually increased over time with levels peaking at 8 weeks after tamoxifen administration (**Fig. 3c**), further demonstrating that meningeal B cells were continuously replenished by HSC-derived progenitors. Similar B cell replenishment dynamics was observed in the BM, whearese the B cell replacement in the spleen, LN and lung was much less efficient within 9 weeks after tamoxifen injection (**Extended Data Fig. 5a**). This is in accordance with the notion that meninges and BM mainly contain early developing B cells with a faster turnover rate from HSCs than mature B cells in other peripheral organs.

**Fig. 3:**
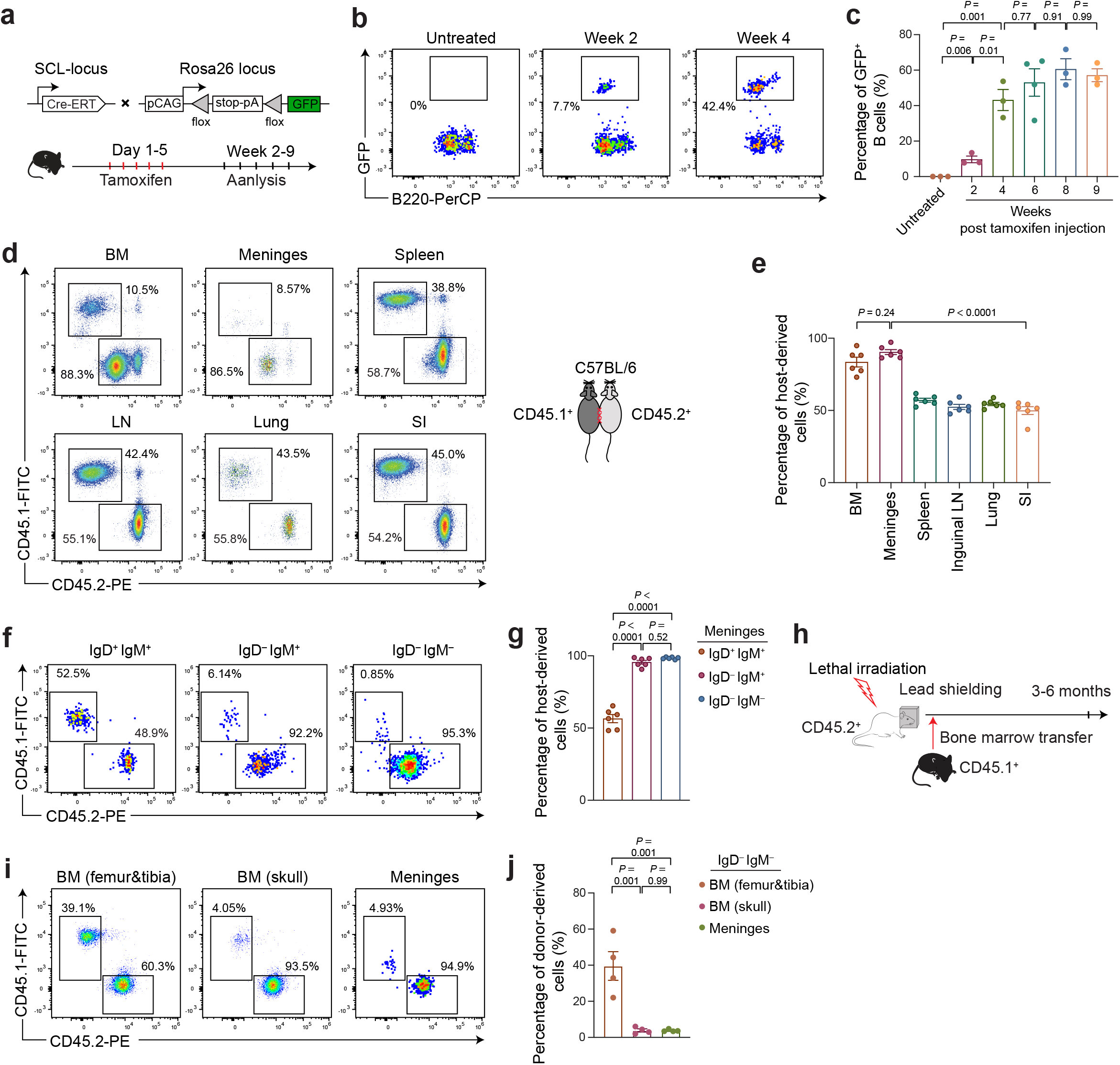
Circulation-independent refreshment of meningeal developing B cells by HSC-derived progenitors. **a**, Schematic illustration of fate mapping of HSC-derived cells by using *SCL-CreERT;R26-GFP* mice. **b,** Representative flow plots showing percentages of GFP^+^ B cells in the meninges at different time points after tamoxifen injection. **c,** Quantification of percentages for GFP^+^ cells in (b). **d,** CD45.1^+^ and CD45.2^+^ mice were surgically connected to generate parabiotic partners (right). Representative flow plots showing percentages of CD45.1^+^ or CD45.2^+^ cells in total B cell (CD19^+^B220^+^) compartment in CD45.2^+^ mice at 8 weeks after surgery (left). **e,** Quantification of percentages for host-derived cells in (d). **f,** Representative flow plots showing percentages of CD45.1^+^ or CD45.2^+^ cells in IgD^+^IgM^+^, IgD^-^IgM^+^ and IgD^-^IgM^-^ B cell compartments in the meninges of CD45.2^+^ mice at 8 weeks after surgery. **g,** Quantification of percentages for host-derived cells in (f). **h**, Experimental design for constructing a chimeric model in which exogenous HSCs engraft less efficiently in the BM of skull compared with femur and tibia. BM cells isolated from CD45.1^+^ mice were transferred to lethally irradiated CD45.2^+^ recipients carrying a lead-cap. Immune cells were isolated and analyzed at 3 to 6 months after the transfer. **i**, Representative flow plots showing percentages of CD45.1^+^ or CD45.2^+^ cells in IgD^-^IgM^-^ B cell compartment in different tissues as indicated. **j,** Quantification of percentages for donor-derived CD45.1^+^ cells in (i). Data are representative of two (b, i) or three (d, f) or the pool from two (c, j) or three (e, g) independent experiments shown as the mean ± s.e.m. Each symbol represents an individual mouse (c, e, g, i). Statistical significance was tested by one-way ANOVA followed by Tukey’s multiple comparisons test (c, e, g, i).

Next, to investigate whether the replenishment of developing B cells in the meninges from HSCs is through circulation, we induced parabiotic mouse model, in which surgically adjoined parabiotic mice can establish a rich anastomotic circulation leading to complete peripheral blood exchange and blood chimerism. We accessed the frequency of partner-derived CD19^+^ B cells, and found a 50:50 exchange rate of B cells in the spleen, LN, lung and SI at 8 weeks after surgery (**Fig. 3d, e**). In striking contrast, total B cells in the meninges and BM from the same animals showed significantly lower exchange rate. Furthermore, wherase the mature B cells (IgD^+^IgM^+^) showed a ~50% exchange rate, the exchange of pre- and pro-B cell (IgD^-^IgM^-^), as well as immature B cells (IgD^-^IgM^+^) was barely detected in the meninges (**Fig. 3f, g**) and BM (**Extended Data Fig. 5b**). Thus, the early developing B cells in the meninges does not rely on the continuous influx of circulating cells from the bloodstream.

The meninges is directly connected with skull marrow cavities via microscopic vascular channels, which serve as a “shortcut” route allowing neutrophils to migrate toward the CNS^19^. In this regard, we then investigated whether HSCs from the skull BM is a preferred origin of developing B cells in the meninges. We transferred BM cells of CD45.1^+^ mice into CD45.2^+^ recipients which carried a protective lead-cap during the pre-irradiation to creat chimaeres (**Fig. 3h**), in which exogenous HSCs less efficiently engrafted in the skull, where HSC-supporting niches were still occupied by the endogenous cells, in compare with bones in the rest of the body, such as femur and tibia (data not shown). In such chimaeres, the BM of skull manifested a distinct cellular composition from that in the femur and tibia, evident by a signigicantly lower frequency of donor-derived cells (CD45.1^+^) in the pro- and pre-B (IgD^-^IgM^-^), as well as immature B (IgD^-^IgM^+^) cell population in the skull BM (**Fig. 3i, j, Extended Data Fig. 5c**). Strikingly, the composition of developing B cells in the meninges was closer to that in the skull rather than femur and tibia, even at 3 to 6 months after the BM transfer (**Fig. 3i, j, Extended Data Fig. 5c**). Given that the turnover time of meningeal B cells were about 6 to 8 weeks (**Extended Data Fig. 5a**), these results demonstrated that HSCs-derived progenitors in the skull appear to be the preferred origin of developing B cells in the meninges. However, it remains to be determined whether HSC-derived progenitors in the skull enter the meninges direcly through the vascular channels^19^.

### Autoreactive B cells recognizing CNS-specific antigens are eliminated in the meninges

The identification of early development B-lineage in the meninges indicates a potential *in situ* tolerance checkpoint for self-reactive B cells recognizing CNS-specific antigens, which otherwise escape negative selection checkpoint in the bone marrow. To test this hypothesis, we leveraged the MOG-specific Ig heavy chain (IgH^MOG^) knock-in (Th) mouse model^5^, in which 40% of B cells in the BM and spleen are reactive to MOG, a self-antigen restricted to the nervous system^5^. We observed a significant reduction of meningeal B cells in Th mice in comparison to wild type (WT). This B cell defect is not simply due to a disruption of B cell development caused by BCR transgene, since the number of total B cells in the meninges remains unaltered in MD4 mice^20^, which carry rearranged heavy and light chain transgenes recognizing hen egg lysozyme (HEL), a nonself-antigen (**Fig. 4a**). Furthermore, the proportion of total IgM^+^ B cells in the meninges of Th mice was significantly decreased compared to the BM in femur and skull, indicating that BCR-expressing B cells were compromised specifically in the meninges (**Fig. 4b, c**).

**Fig. 4:**
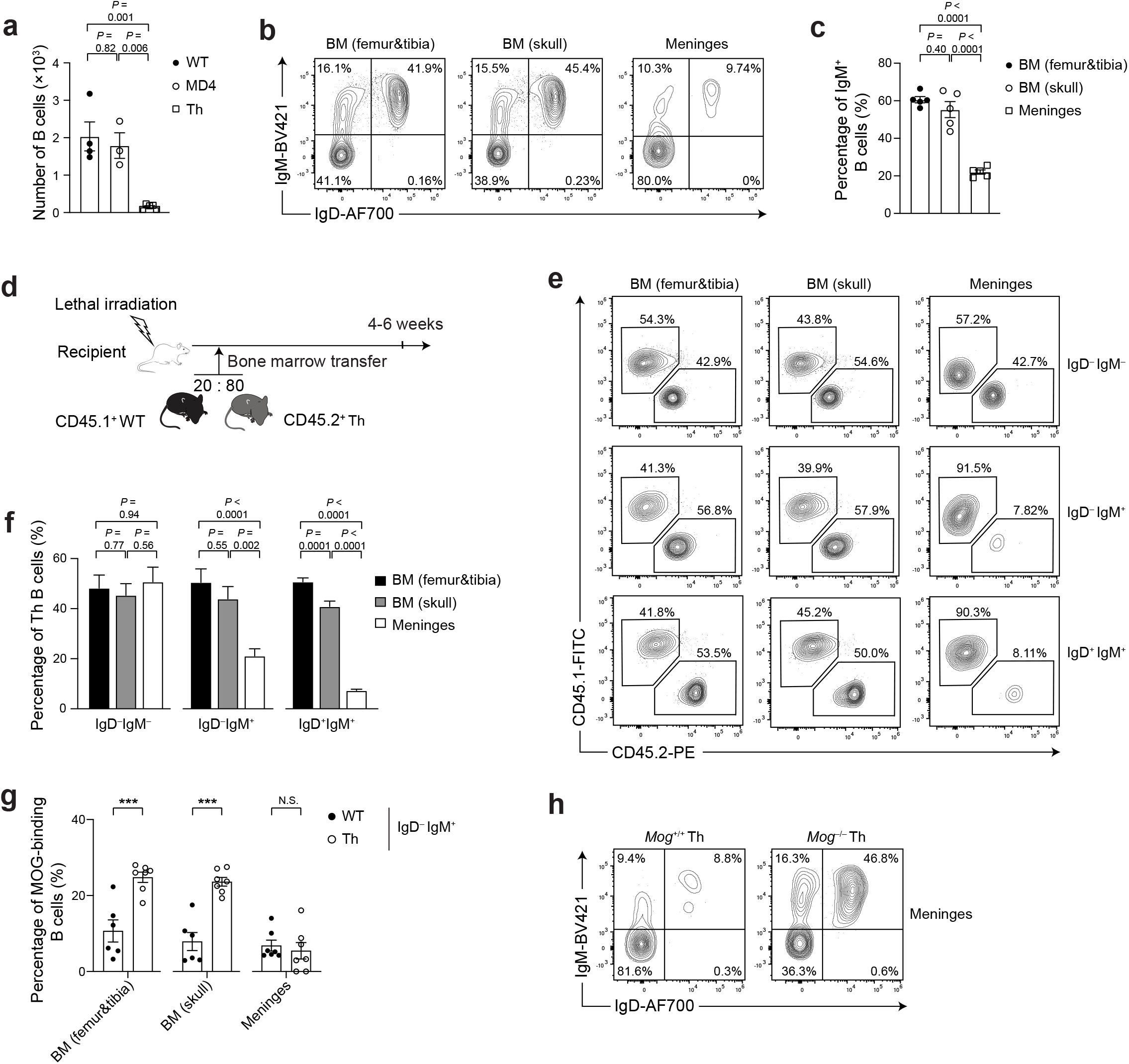
CNS antigen-reactive B cells are eliminated in the meninges but not BM. **a**, Quantification of B cell (B220^+^ CD19^+^) numbers in the meninges of WT, MD4 and Th mice. **b**, Representative flow plots showing membrane IgD and IgM expression in BM and meningeal B cells. **c**, Quantification of percentages for IgD^-^IgM^-^ B cells in (b). **d**, Experimental design of BM chimeric model. **e**, Representative flow plots showing percentages of CD45.1^+^ WT or CD45.2^+^ Th B cells in IgD^-^IgM^-^, IgD^-^IgM^+^ and IgD^+^IgM^+^ B cell compartments in the BM and meninges. **f,** Quantification of percentages for CD45.2^+^ Th cells in (e). **g,** Quantification of percentages for MOG-binding cells in the IgD^-^IgM^+^ B cell compartment in **Extended Data Fig. 6a-c**. **h**, Representative flow plots showing membrane IgD and IgM expression in meningeal B cells of *Mog*^+/+^ (left) and *Mog*^-/-^ (right) Th mice. Data are representative of two (b,h) or five (e) or the pool from two (a, c) or five (f) or three (g) independent experiments shown as the mean ± s.e.m. Each symbol represents an individual mouse (a, c, g). Statistical significance was tested by one-way ANOVA followed by Tukey’s multiple comparisons test (a, c, f) or two-tailed t-test (g).

To further dissect whether the reduction of B cells in the meninges of Th mice is a BCR-driven process, we generated chimeras by transferring bone marrow cells from CD45.1^+^ WT and CD45.2^+^ Th mice at a 20:80 ratio into pre-irradiated WT recipients (**Fig. 4d**). A successful chimerization was confirmed by a 1:4 ratio of CD45.1^+^ and CD45.2^+^ non-B cell populations in the BM and spleen at 4-6 weeks after transplantation (data not shown). We observed a selective defect of Th-derived cells in the immature (IgD^-^IgM^+^) and mature (IgD^+^IgM^+^) B cell compartment within the meninges, but not in the pre- and pro-B (IgD^-^IgM^-^) cell fractions, which exihited similar frequencies of Th-derived B cells as in the femur and skull BM (**Fig. 4e, f**). These observations indicate a specific BCR-dependent reduction of Th-derived B cells in the meninges.

In the BM, autoreactive B cell progenitors recognizing self antigen will be eliminated or inactivated through an antigen-dependent central tolerance mechanism. In this regard, we next directly tested whether MOG-binding competence of B cells was compromised in the meninges of Th mice. In accordance with previous reports^5,21^, the frequencies of immature and mature B cells bound by recombinant His-tagged MOG protein in the BM of skull and tibia of Th mice were significantly increased in compare with WT controls (**Fig. 4g, Extended Data Fig. 6a-d**). Strikingly, the frequencies of MOG-binding B cells in the meninges of Th mice declined to the background level as in WT controls (**Fig. 4g, Extended Data Fig. 6a-d**), demonstrating that MOG-specific B cells were specifically eliminated from the meninges but not BM of skull and tibia. Importantly, this effect relies on the endogeneous expression of MOG, as the genetic ablation of MOG fully restored the frequency of IgM^+^ meningeal B cells in Th mice (**Fig. 4h**). Overall, we conclude that autoreactive B cells recognizing CNS-specific antigens are eliminated in an autoantigen-dependent manner in the meninges.

Our findings discovered a consecutive trajectory of early developing B cells within the meninges of mice and nonhuman primates. Importantly, we revealed that meningeal immature B cells undergo a CNS antigen-dependent tolerance checkpoint to avoid local presence of autoreactive mature B cells, which could contribute to the development of neurological diseases. Intriguingly, as mature B cells were exchangeable between the meninges and periphery through the circulation, the complete absence of MOG-binding mature B cells in the meninges imply that peripheral MOG-specific mature B cells, which escaped the tolerance checkpoints in the BM and spleen, could be also depleted if they reach the meninges. The similar observation of eliminating autoreactive mature B cells was previously observed in the liver of 3–83 BCR transgenic mouse^22^. Therefore, the emergence of *in situ* early developing B cells could function as an additional layer of checkpoint mechanism that stringently restain a local non-self-reactive immune repertoire in the meninges. However, it remains to be elucidated whether early B cell development can also regulate meningeal immunity via other mechanism(s). Local perturbation of B cell development and selection specifically in the meninges is needed to further explore the physiological function and destination of meningeal developing B cells.

## Acknowledgments

We thank H. Wekerle and G. Krishnamoorthy for providing Th mice; J. Zheng for providing Scl-creERT mice; S.M. Kerfoot for sharing mMOGtag plasmid; P. Lu for MOG protein expression and purification; W. Zhu for mouse genotyping; all the members of the Xu and He laboratories for discussion and suggestions; the Laboratory Animal Resources Center, High-Performance Computing Center, Flow Cytometry Core and Genomic Core in Westlake University for their support. This work was supported by the National Key R&D program of China (grants 2020YFA0804200 and 2019YFA0802900), National Natural Science Foundation of China (grants 31970842, U20A20346 and 32070953), and the Education Foundation of Westlake University.

## Author Contributions

D.H. and H.X. conceived and supervised this study. D.C. performed computational analysis; Y.W. performed the experiments with help from D.X. and C.H.; H.X. and D.H. wrote the manuscript with input from all authors.

---

**Extended Data Fig. 1:**
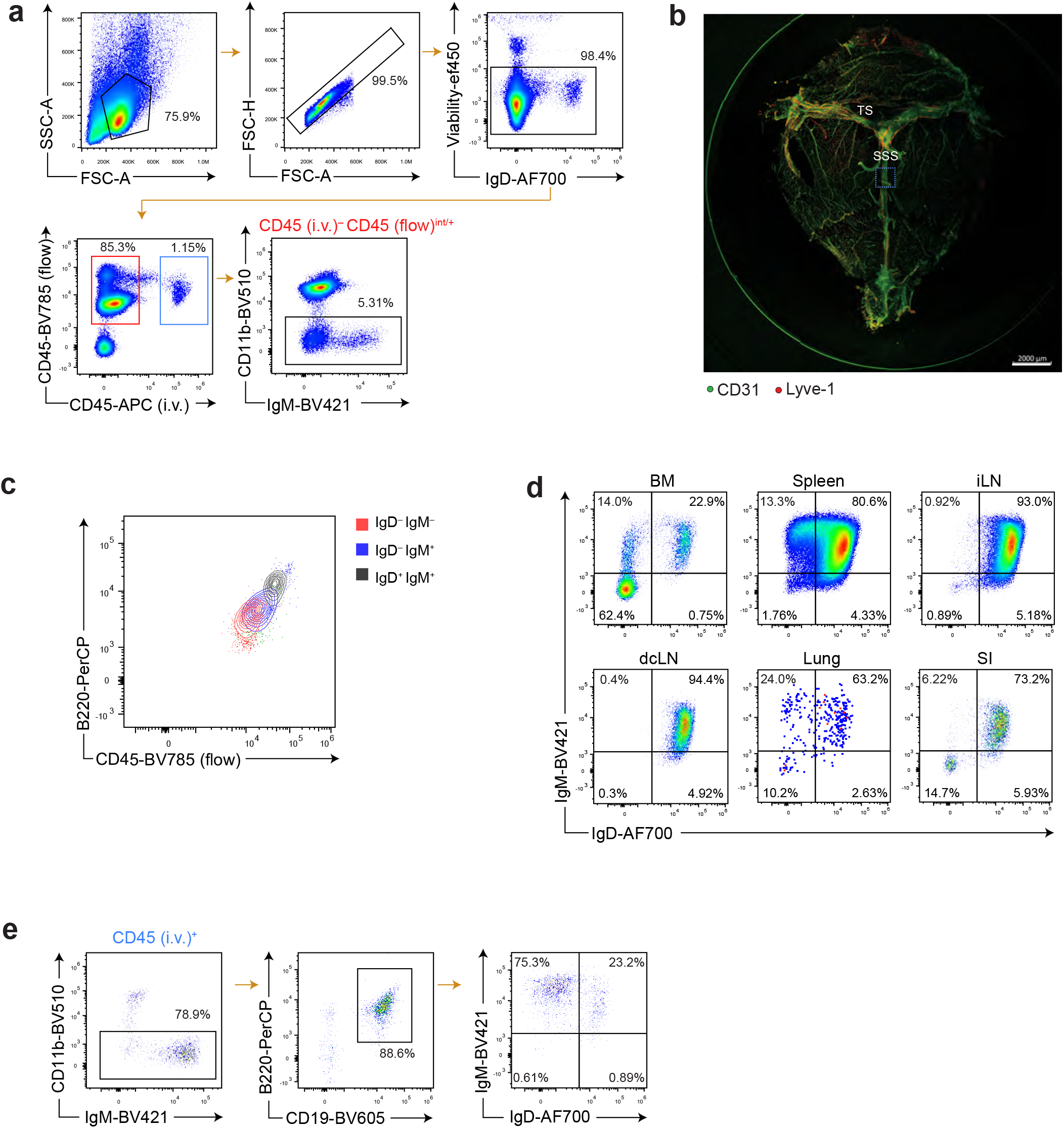
Flow and imaging analysis for meningeal B cells. **a**, Gating strategy for identifying CD45^int/+^CD11b^-^ immune cell population that weren’t labeled by intravenously injected CD45 antibody. **b**, A representative image of CD31 and Lyve-1 labelling in whole-mount meninges. Box: the area for higher magnification imaging in **Fig. 1c**. SSS, superior sagittal sinus; TS, transverse sinus. **c**, Representative FACS plots showing B220 and CD45 expression in IgD^-^IgM^-^ (red), IgD^-^IgM^+^ (blue) and IgD^+^IgM^+^ (black) meningeal B cells. **d**, Representative flow plots showing membrane IgD and IgM expression in B cells (CD19^+^B220^+^) of different tissues as indicated. **e**, Representative flow plots showing membrane IgD and IgM expression in B cells that were labeled by intravenously injected CD45 antibody (blue box, **Extended Data Fig. 1a**). Data are representative of three independent experiments (a, c-e) or biological replicates (b).

**Extended Data Fig. 2:**
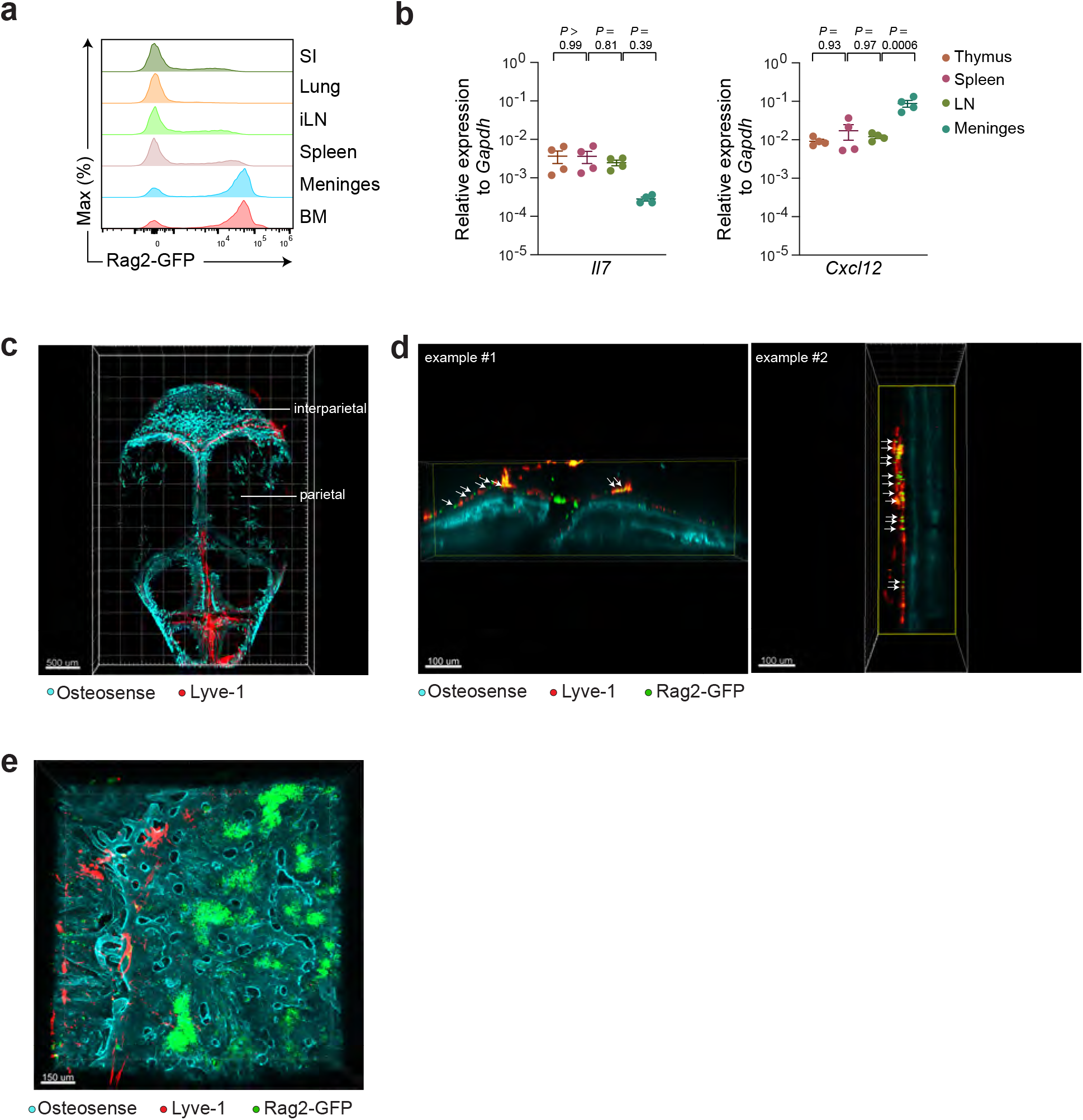
Flow cytometry and image analysis of RAG2-expressing B cells, and real-time PCR quantification of *Il7* and *Cxcl2* expression. **a**, Representative offset flow histograms showing Rag2-GFP expression in B cells isolated from the indicated tissues. **b**, Expression of *Il7* (left) and *Cxcl12* (right) in the thymus, spleen, LN and meninges of WT mice. **c**, A representative three-dimensional (3D) image of a whole mouse skull adhered with dura matter after tissue clearing. The interparietal and parietal area are labeled. Bone outline is visualized with Osteosense. **d**, Representative optical slides of high magnification 3D images near the SSS region in the parietal area of skull. Arrow, RAG2-GFP-expressing cells align linearly within Lyve1-expressing lymphatic vessels. **e**, A representative high magnification 3D image of interparietal area of skull. Data are representative of three (a) or two (c-d) biological replicates or the pool of four biological replicates shown as the mean ± s.e.m (b). Each symbol represents an individual mouse and statistical significance was tested by one-way ANOVA followed by Tukey’s multiple comparisons test (b).

**Extended Data Fig. 3:**
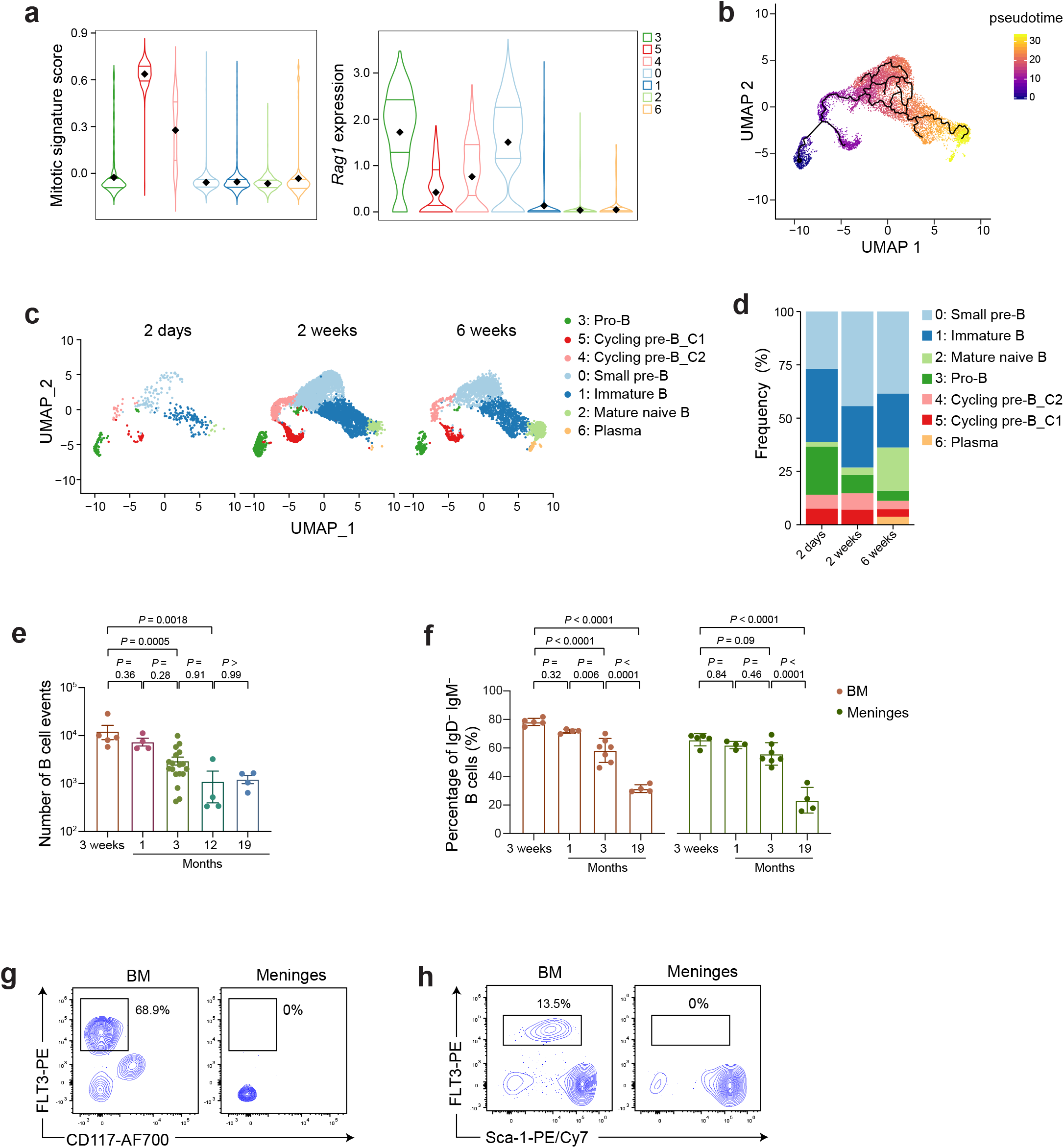
scRNA-seq and flow cytometry analysis of meningeal B cells. **a**, Violin plots of the distribution of mitotic signature scores (**Online Methods**) and *Rag1* gene expression in scRNA-seq data of cells in cluster 0 through 6. Diamond indicates the mean; lines, first and third quartiles. **b**, Pseudotime trajectory of GC B cells constructed by Monocle. **c**, Shown are 2D embedding as in **Fig. 1g**, separated according to different ages of mice. **d**, Distribution of proportions for each B cell clusters at the different ages in (c). **e**, Quantification of numbers of B cells in the meninges at different ages as indicated. **f**, Quantification of frequencies of IgD^-^IgM^-^ cells in total B cell population (B220^+^CD19^+^) in the meninges and BM at different ages as indicated. **g**, Representative flow plots of pre-pro B cells in the BM and meninges. The cells were gated on CD45^+/int^Ly6C^-^CD3^-^CD11b^-^GR1^-^CD19^-^B220^+^CD24^low^CD43^+^ population. **h**, Representative flow plots of CLPs in the BM and meninges. The cells were gated on CD45^+/int^Lin^-^CD127^+^population. Data are the pool from six biological replicates (a-d) or representative of three (g, h) biological replicates or the pool from three independent experiments (e,f) shown as the mean ± s.e.m. Each symbol represents an individual mouse (e,f). Statistical significance was tested by one-way ANOVA followed by Tukey’s multiple comparisons test (e,f).

**Extended Data Fig. 4:**
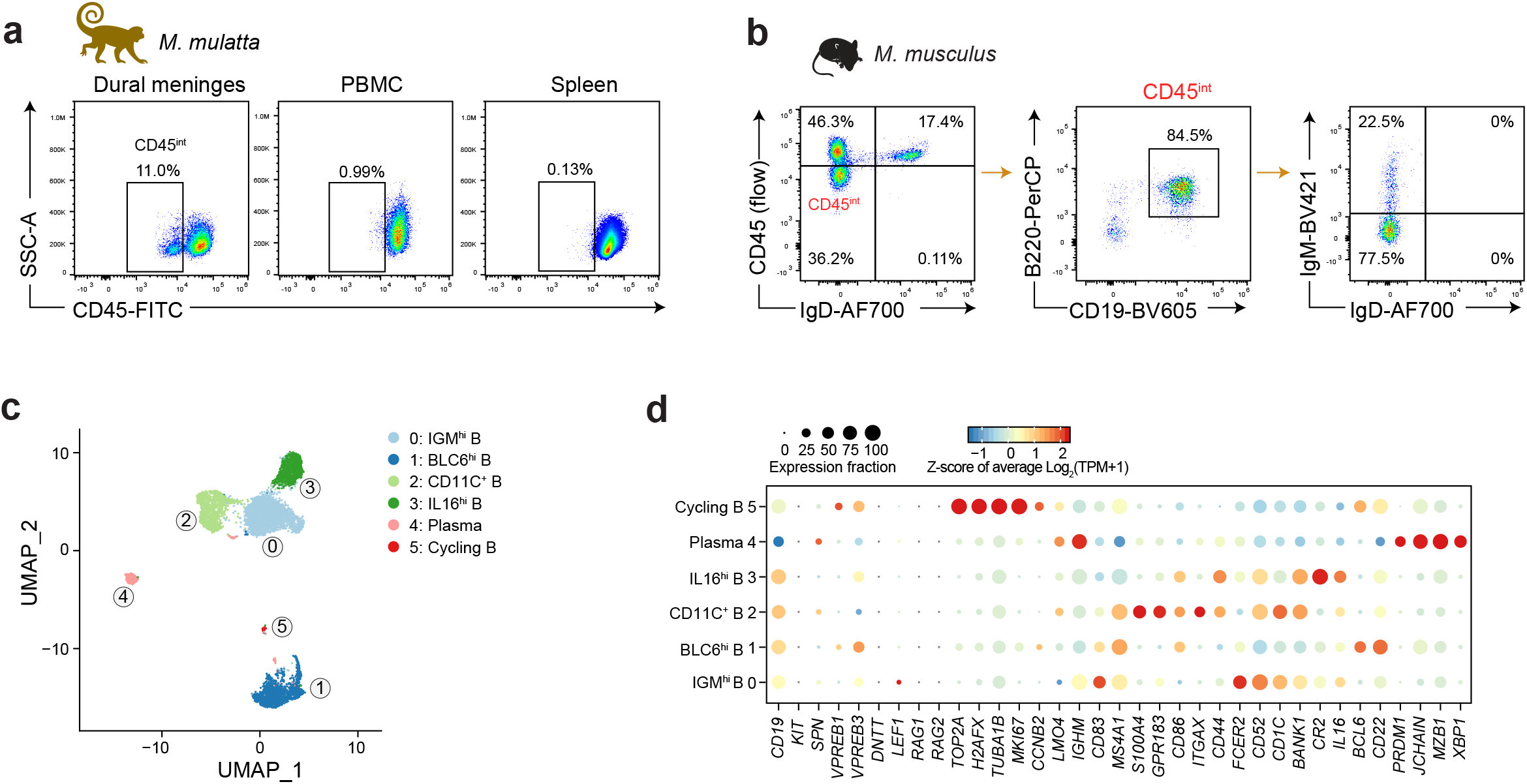
scRNA-seq and flow cytometry analysis of NHP B cells. **a**, Representative FACS plots of CD45^int^ cells in the dural meninges, PBMC and spleen of NHPs. **b**, Representative FACS plots showing that CD45^int^CD19^+^B220^+^ cells were at the early developing stage lacking IgD or both IgD and IgM expression in mice. **c**, UMAP learned 2D representation of spleen B cell profiles (dots) of NHPs, colored and numbered by cluster membership. Clusters are rank ordered by size. **d**, For representative differentially expressed genes (columns) across clusters (rows) in (c), shown is the fraction of cells in the cluster that express a gene (dot size) and the z-score of the mean expression of that gene in the cluster (color; z-score of average log2(TPM+1)). The inferred identities were labeled at the left. Data are representative of three (a,b) or the pool of two (c,d) replicates.

**Extended Data Fig. 5:**
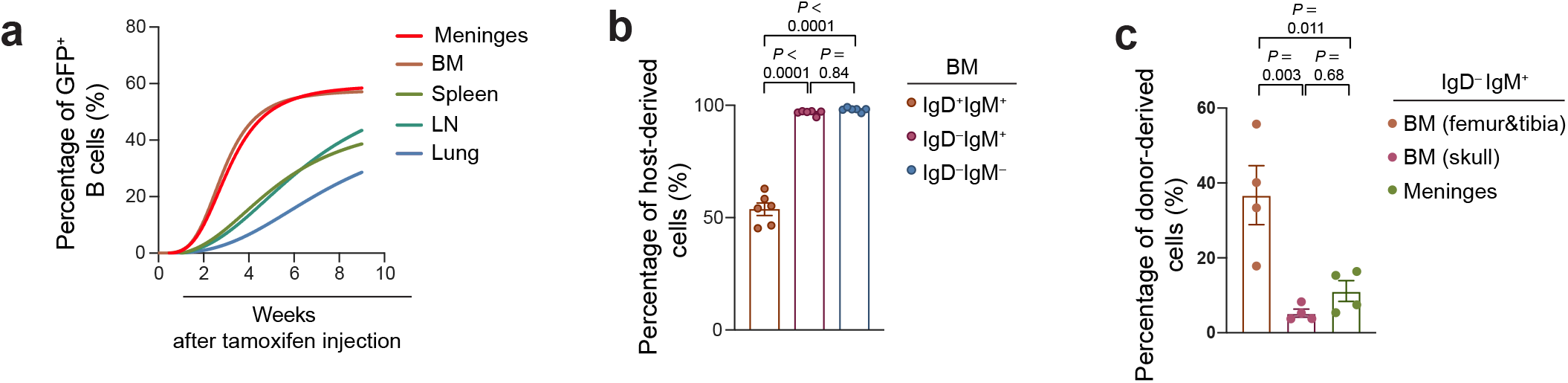
Quantification of frequencies of B cells in the lineage-tracing and BM chimeric models. **a**, Shown are nonlinear regression curve fitting percentages of GFP^+^ B cells over time in the indicated tissues. **b,** Quantification of percentages for host-derived cells in IgD^+^IgM^+^, IgD^-^IgM^+^ and IgD^-^IgM^-^ B cell compartments in the BM at 8 weeks after parabiotic surgery as in **Fig. 3d**. **c,** Quantification of percentages for donor-derived cells in IgD^-^IgM^+^ B cell compartment in the meninges at 3-6 months after the BM transfer as in **Fig. 3h**. Data are the pool from two (a, c) or three (b) independent experiments shown as the mean ± s.e.m. Each symbol represents an individual mouse (b, c). Statistical significance was tested by one-way ANOVA followed by Tukey’s multiple comparisons test (b, c).

**Extended Data Fig. 6:**
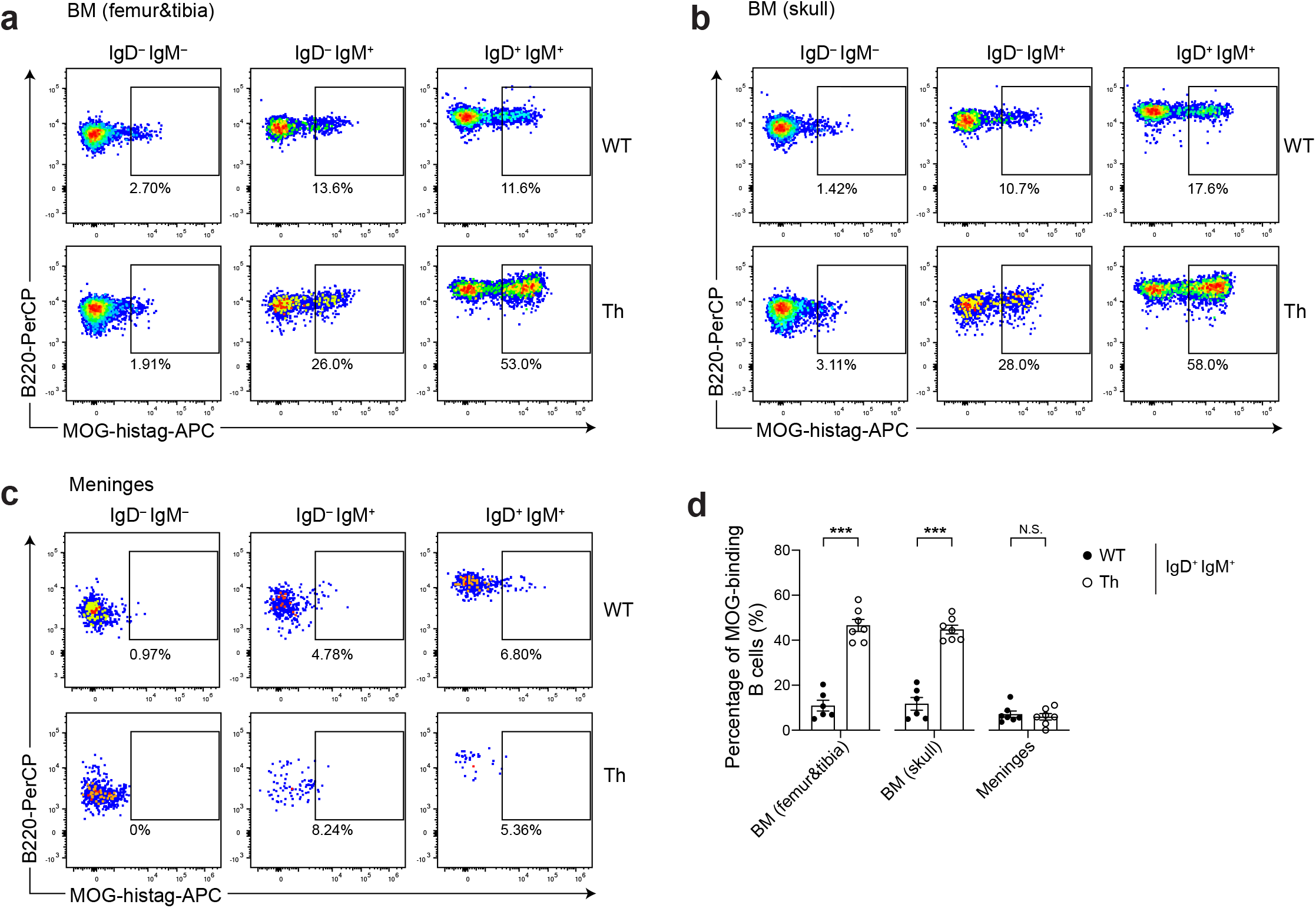
Flow cytometry analysis of MOG-binding B cells in the BM and meninges of WT and Th mice. **a-c**, Representative flow plots showing percentages of MOG-binding cells in IgD^-^IgM^-^, IgD^-^ IgM^+^ and IgD^+^IgM^+^ B cell compartments in the femur (a) or skull (b) BM or meninges (c) of WT and Th mice. **d**, Quantification of percentages for MOG-binding cells in the IgD^+^IgM^+^ B cell compartment in (a-c). Data are representative (a-c) or the pool (d) of three independent experiments shown as the mean ± s.e.m. Each symbol represents an individual mouse (d). Statistical significance was tested by two-tailed t-test (d).

**Supplementary movie 1:** Large volumetric C_e_3D microscopy of the skull attached with dura matter. Bone outline is visualized with Osteosense (cyan). The tissues were stained with Lyve-1 antibody (red), cleared, and imaged via confocal microscopy.

**Supplementary movie 2:** Volumetric C_e_3D imaging of parietal area of skull of RAG2-GFP mice (Cyan: Osteosense, Red: Lyve-1, Green: RAG2-GFP).

**Supplementary movie 3:** Volumetric Ce3D imaging of interparietal area of skull of RAG2-GFP mice (Cyan: Osteosense, Red: Lyve-1, Green: RAG2-GFP).

**Supplementary Table 1:** Differential gene expression analysis between cells in each cluster and the remaining cells in mice meningeal B cell dataset

**Supplementary Table 2:** Differential gene expression analysis between cells in each cluster and the remaining cells in NHP meningeal B cell dataset

**Supplementary Table 3:** Differential gene expression analysis between cells in each cluster and the remaining cells in NHP spleen B cell dataset

**Supplementary Table 4.** Reagents used in this study

## Methods

### Animals

C56BL/6J (Jax 000664), CD45.1 (Jax 002014), μMT (Jax 002288), MD4 (Jax 002595) and Rag2-EGFP (005688) were originally from the Jackson Laboratory. *Mog*^-/-^ (NM-KO-190118) mice were purchased from Shanghai Model Organisms Center. Th mice were generously provided by H. Wekerle and G. Krishnamoorthy (Max Planck Institute of Neurobiology). HSC-Scl-CreERT mice and Ai47 GFP reporter mice were kindly provided by J. Zheng (Shanghai Jiao Tong University) and Z. Qiu (Chinese Academy of Sciences), respectively. Rag2-EGFP mice were bred to C57BL/6J mice for at least 10 generations. HSC-Scl-CreERT and Ai47 mice were crossed to obtain *Scl-Cre* × *Ai47* strain, *Mog*^-/-^ and Th mice were crossed to obtain *Mog*^-/-^ × *Th* strain. Mice were housed in specific-pathogen-free conditions at ~22°C with humidity of 40-70% and a light/dark cycle of 12 hours, and were used and maintained in accordance with the Institutional Animal Care and Use Committee guidelines of Westlake University. Both male and female mice were used at 8–18 weeks unless otherwise indicated. Tamoxifen was dissolved in corn oil at 37□°C with shaking overnight to a working concentration of 20 mg/ml and was administered intraperitoneally at a dose of 75 mg/kg when indicated.

*Macaca mulatta* used in this study were maintained at ~25°C on a light/dark cycle of 12 hours and raised at Xieerxin Biology Resource with accreditation of Laboratory Animal Care accredited facility in Beijing, in compliance with all local and federal laws governing animal research. None of the animals used in this study had a clinical or experimental history. PBMC, spleen and dura mater from 3-8 years old animals were collected. This study was conducted in accordance with the Principles for the Ethical Treatment of Non-Human Primates and was approved by the Institutional Animal Care and Use Committee of Westlake University.

### Parabiosis

Two six weeks old female mice (CD45.1 and CD45.2) of similar size were co-housed for 2 weeks prior to the procedure. The mice were anesthetized with i.p. injection of avertin to full muscle relaxation for surgery. The corresponding lateral aspect of each mouse was shaved, and an incision was made from the olecranon to the knee joint of each mouse, bluntly dissecting the subcutaneous fascia to create 0.5□cm of free skin on each side of the incision. The olecranon and knee joints were attached by an absorbable 5-0 suture. The dermises of the parabiotic partners were pushed together (excluding epidermal layers in the junction of the dermis) and closed with sutures. Meloxicam was administrated right after surgery for pain control and repeated once in 24 hours. Parabiotic mice were maintained under antibiotic-containing water (0.8 g/L Sulfamethoxazole and 0.4 g/L Trimethoprim) for 2 weeks after surgery and then returned to regular drinking water.

### Bone marrow transfer model

CD45.2 B6 recipients were lethally irradiated by X-ray (5□Gy□×□2), and then intravenously transferred with a combination of 4□×□10^6^ bone-marrow leukocytes from indicated donors mixed according to the indicated ratio. When indicated, the recipients carried lead cap during the irradiation. Chimeric mice were maintained under antibiotic-containing water ( 0.8 g/L Sulfamethoxazole and 0.4 g/L Trimethoprim) for 3-4 weeks after reconstitution and then returned to regular drinking water.

### Tissue preparation and immune cell isolation

After intravenous injection of APC-conjugated CD45 antibody, mice were anesthetized with intraperitoneal injection of avertin, followed by transcardially perfused with 20 mL of icecold PBS. Skin was removed from the head and the muscle was stripped from the bone. The top of the skull was collected, and dura mater was carefully removed from the skull using fine tissue forceps under a stereo microscope. Lung, brain and dura mater were chopped and subject to enzymatic digestion in Hanks’ Balanced Salt Solution (HBSS) containing 2.5 mg/mL collagenase D (Roche) and 0.1 mg/mL DNase I (Roche) for 25 minutes at 37°C with gentle shaking every 5 minutes. The supernatants were then passed through 100μm filters. After washing once with MACS buffer (1×PBS pH 7.4; PBS plus 2% FCS and 2 mM EDTA), cells were resuspended in 37% percoll and centrifuged for 20 minutes at 600×g at 25 °C. Splenocytes and bone marrow cells were mechanically harvested. Erythrocytes were depleted using ACK lysis buffer.

For isolating immune cells from the dura matter of NHP, animals were anesthetized and perfused with 2L ice-cold PBS with heparin. After opening the top of skull, the dura mater was carefully removed from the brain and then chopped for enzymatic digestion in Hanks’ Balanced Salt Solution (HBSS) containing 15 mg/mL collagenase D (Roche) and 1 mg/mL DNase (Roche) for 1h at 37 °C with gentle shaking every 5 minutes. The supernatants were then processed as described above for murine cells.

### Flow cytometry and cell sorting

Cells were washed and prepared as single-cell suspensions in MACS buffer (1×PBS containing 2% FCS and 10 mM EDTA). Nonspecific antibody binding was blocked with Fc-blocking antibody 2.4G2 (BD, 25 μg/ml) by incubation for 10 min on ice. Cells were stained for 30 min at 4 °C with indicated antibody cocktails. For labeling mog-specific B cells, splenocytes were incubated with purified His-tagged MOG protein (20μg/ml) for 30 minutes, followed by incubation with AF647 conjugated anti-His-tag antibody for 30 minutes at 4 °C after washing twice. All antibodies and relevant reagents used for flow cytometry are listed in Supplementary Table 4. Data were collected on Cytoflex (Beckman Coulter) and analyzed using FlowJo 10.7.1 software (TreeStar). Murine CD45^+^CD11b^-^ and macaque CD45^+^CD11b^-^CD3^-^ meningeal cells were sorted using MA900 (Sony) with a 100-μm chip at 4 °C.

### Meninges immunohistochemistry

After anesthetization and transcardial perfusion, the skull cap adhered with intact dura mater was harvested and fixed in 4% paraformaldehyde for 16–18 hours at 4°C. After washing three times in PBS for 5 minutes, whole-mount meninges were peeled off from the skull cap for immunostaining. In brief, whole-mount meninges were blocked and permeabilized in staining buffer (1×PBS containing 0.2% Triton X-100, 0.05% Tween and 2% BSA) for 1□hour at 25 °C. Meninges were then incubated with anti-Lyve-1 (1:400) and anti-B220 (1:400) (Supplementary Table 4) in the staining buffer at 4□°C for 24□hours. Whole mounts were then washed 3 times for 5 minutes at 25 °C in 0.1 M Tris-HCl followed by incubation with Cy3 conjugated anti-rabbit IgG antibody (1:1000 dilution) for 2□hours at 25 °C with gentle agitation. Sections were washed once in 0.1 M Tris-HCl and incubated with 4’,6-diamidino-2-phenylindole (DAPI; 0.3□μg/ml) in 0.1 M Tris-HCl. Using fine surgical forceps, the dura mater was flattened on a glass slide and mounted. After 5 min in DAPI reagent (0.3□μg/ml), a whole mount tissue was flattened and attached on a glass slide and mounted with ProlongGold Antifade reagent (Thermo). Images were acquired with Zeiss LSM800 confocal system. For imaging complete whole mount meninges, images were acquired with a 2.5× objective with 0.12 NA. Other confocal images were acquired using a 20× objective with 0.8 NA. The results were analyzed with Imaris (Bitplane) software.

### C_e_3D skull clearing and imaging

RAG2-GFP mice were intravenously injected with OsteoSense 680 EX (2nmol in 100 μl PBS, PerkinElmer) for *in vivo* labeling bone structure, followed by perfusion with PBS and later 4% paraformaldehyde (PFA). Then the skull cap adhered with intact dura mater were harvested and subjected to tissue clearing using Ce3D protocol as previously reported^14^. In brief, the skull was fixed in PLP buffer (0.075 M lysine, 0.37 M sodium phosphate (pH 7.2), 1% formaldehyde, and 0.01 M NaIO4) at 4°C overnight. After washing three times in washing buffer (1×PBS containing 0.2% (v/v) Triton X-100 and 0.5% (v/v) 1-thioglycerol) for 30-60 minutes, the skull was incubated in blocking buffer (1XPBS containing 1% (v/v) BSA, 1% (v/v) normal mouse serum and 0.3% (v/v) Triton X-100) at 37°C for 24 hours in a shaker (200 rpm). Then the tissue was incubated in the blocking buffer containing anti-Lyve-1 antibody (1:400) (Supplementary Table 4) for 1 day at 37°C in a shaker (200 rpm), followed by incubation with the fresh washing buffer for 14 hours at 37°C and then 1 day at 25 °C. The skull was then briefly dried on a Kimwipe and immediately immersed in Ce3D clearing solution (1× PBS containing 22% (v/v) N-methylacetamide, 0.8 g/ml Histodenz, 0.1% (v/v) Triton X-100 and 0.5% (v/v) 1-thioglycerol) until clear enough for imaging. All procedures should be protected from light.

### Real-time PCR

Total RNA was extracted with Trizol (ThermoFisher Scientific) and cDNA was synthesized using iScript cDNA Synthesis Kit (Biorad) according to the manufacturer’s protocols. Quantitative PCR was performed on a Qtower384G System (Jena) with iTaq Universal SYBR Green Supermix (BioRad). Primers used for quantitative PCR are in Supplementary Table 4.

### Droplet-based scRNA-seq

Single cells were captured via the GemCode Single Cell Platform using the GemCode Gel Bead, Chip and 3□-end Library Kits (10X Genomics), according to the manufacturer’s protocol. Briefly, flow-sorted murine meningeal cells (CD45^-^CD11b^-^), which were prelabeled with hashtag antibodies (Biolegend), were pooled and loaded at 30,000 cells per channel. Flow-sorted macaque immune cell population (CD45^-^CD11b^-^CD3^-^) were loaded at 8,000 cells per channel. The cells were then partitioned into the GemCode instrument, where individual cells were lysed and mixed with beads carrying unique barcodes in individual oil droplets. The products were subjected to reverse transcription, emulsion breaking, cDNA amplification, and sample index attachment. Libraries were pair-end (150+ 150 bp) sequenced on a NovaSeq (Illumina).

### scRNA-seq data processing

For murine dataset, initial processing and gene expression estimation were performed using cellranger count (v3.1.0) with the refdata-cellranger-mm10-3.0.0 reference from 10X Genomics for alignments. The unique molecular identifier (UMI) count matrix was converted to Seurat objects using the R package Seurat (v3.1.0) operated in RStudio (v1.4). Cells from different time points and animals were demultiplexed based on hashtag using *HTODemux* function with default parameters. The cells with more than two or without tags were removed. After further removing low-quality cells with less than 1200 genes, more than 20000 UMIs or more than 10% of UMIs mapped to mitochondrial genes, we obtained 12324 cells (403, 3106, 2966, 3079 and 2770 cells from 2-day, 2-week, 2-week, 6-week and 6-week mice, respectively). 17508 genes were retained after filtering genes expressed in less than 3 cells. The filtered gene expression matrix was normalized using Seurat’s *NormalizeData* function and 2000 highly variable genes were identified per dataset using Seurat’s *FindVariableFeatures* function. After we scaled the normalized data, highly variable genes were selected for principal component analysis (PCA). The first 20 principal components (PCs) were used for Uniform Manifold Approximation and Projection (UMAP) for visualization and graph-based clustering with resolution set to 0.3. Using normalized expression values, marker genes for each cluster were inferred by the MAST test as implemented in the *FindAllMarker* function of Seurat. According to the marker genes identified, B cell and plasma cell clusters were subset for downstream analysis. To remove effects from cell cycle, we used *CellCycleScoring* function to assign each cell a score based on its expression of G1/S and G2/M markers, and then regressed out it during data scaling. After that, we reran the PCA and used first 15 PCs for UMAP and clustering. Marker genes for new clusters were also identified by the MAST test.

For NHP dataset, we firstly built a custom reference genome of *Macaca mulatta* according to 10X Genomics’s tutorial. Genome FASTA file (Macaca mulatta.Mmul_10.dna.toplevel.fa) and GTF file (Macaca_mulatta.Mmul_10.102.chr.gtf) were download from Ensembl. Entries in GTF file for non-polyA transcripts that overlap with protein-coding gene models were filtered using cellranger mkgtf (v3.1.0), and then the reference were created using cellranger mkref. For meningeal samples, cells with less than 800 genes, more than 12000 UMIs or more than 5% of UMIs mapped to mitochondrial genes were removed. We got 7051 cells (3457 and 3594 from replicates 1 and 2, separately). 14507 genes were retained after filtering genes expressed in less than 3 cells. For spleenocytes, cells with less than 900 genes, more than 5% of UMIs mapped to mitochondrial genes, less than 2000 UMIs or more than 10000 UMIs were removed. The filtered gene expression matrix was then normalized, clustered, and visualized as above in murine dataset.

### Pseudotime Inference

Differentiation trajectories for B cell clusters were constructed using Monocle 3 (v0.2.0). The *cell_data_set* object used for Monocle 3 were constructed based on Seurat processing as mentioned before. Then, we fit a principal graph using the *learn_graph* function and chosen the root nodes manually to order the cells according to their progress through the developmental program.

### Gene signature score

Gene signature scores were calculated as previously reported^15^. Briefly, for each targeted gene set, we chose 10 times more genes as a control gene set, which were defined by finding the closest genes in terms of expression level and detection rate, aiming to has a comparable distribution. As the control gene set is 10-fold larger, such that its average expression (log2 (CPM+1)) is analogous to averaging over 10 randomly selected gene sets of the same size as the targeted gene set. We then obtained a gene signature score by subtracting average expression value of a control gene set from average expression value of the targeted gene set for individual cells.

### Statistical analysis

Prism 8 (GraphPad Software) was used to perform one-way ANOVA followed by Tukey’s multiple comparisons test and two-tailed *t*-test (except for RNA-seq data for which we use R (v3.6.1) for statistical analysis).

### Data Availability

We’re depositing scRNA-Seq data into the Gene Expression Omnibus database. The accession number will be provided when it’s available.

### Code availability

Code used in this study are available from the corresponding authors upon reasonable.

